# Natural variation further increases resilience of sorghum bred for chronically drought-prone environments

**DOI:** 10.1101/2021.10.20.465110

**Authors:** Hongxu Dong, Techale Birhan, Nezif Abajebel, Misganu Wakjira, Tesfaye Mitiku, Cornelia Lemke, Vincent Vadez, Andrew H. Paterson, Kassahun Bantte

## Abstract

Climate-change-associated shifts in rainfall distribution together with a looming worldwide water crisis make drought resilience of central importance to food security. Even for relatively drought resilient crops such as sorghum, moisture stress is nonetheless one of the major constraints for production. Here, we explore the potential to use natural genetic variation to build on the inherent drought tolerance of an elite cultivar (Teshale) bred for Ethiopian conditions including chronic drought, evaluating a backcross nested-association mapping (BC-NAM) population using 12 diverse founder lines crossed with Teshale under three drought-prone environments in Ethiopia. All twelve populations averaged higher head exsertion and lower leaf senescence than the recurrent parent in the two highest-stress environments, reflecting new drought resilience mechanisms from the donors. A total of 154 QTLs were detected for eight drought responsive traits – the validity of these were supported in that 100 (64.9%) overlapped with QTLs previously detected for the same traits, concentrated in regions previously associated with ‘stay-green’ traits as well as the flowering regulator *Ma6* and drought resistant gene *P5CS2*. Allele effects show that some favorable alleles are already present in the Ethiopian cultivar, however the exotic donors offer rich scope for increasing drought resilience. Using model-selected SNPs associated with eight traits in this study and three in a companion study, phenotypic prediction accuracies for grain yield were equivalent to genome-wide SNPs and were significantly better than random SNPs, indicating that these studied traits are predictive of sorghum grain yield. Rich scope for improving drought resilience even in cultivars bred for drought-prone regions, together with phenotypic prediction accuracy for grain yield, provides a foundation to enhance food security in drought-prone areas like the African Sahel.

## Introduction

In drier parts of the world, more than a half-century of climate change coupled with ongoing population growth has already pushed agriculture increasingly into ecologically marginal regions and is expected to severely impact agricultural production especially in sub-Saharan Africa (Kurukulasuriya et al., 2006; Pironon et al., 2019). In the African Sahel, years of above-average rainfall followed by drought starting in the 1960’s (Haywood, Jones, Bellouin, & Stephenson, 2013), were closely associated with more than doubling of the area devoted to crops (Kandji, Verchot, & Mackensen, 2006). Such unsustainable land management practices are well known to deplete topsoil (Desta, Tamene, Abera, Amede, & Whitbread, 2021) and contribute to the Sahel ranking among the regions with most severe soil degradation worldwide (ISRIC, 1990). Indeed, depleting soil organic matter not only increases atmospheric carbon but also sacrifices the potential of soil as a carbon reservoir (Amelung et al., 2020).

Its unusual tolerance of low inputs, especially water, make the cereal crop sorghum essential in rain-fed low-input cropping regions that were inadequately addressed in the ‘Green Revolution’ (Cantrell & Hettel, 2004; Conway, 1997; Paterson & Li, 2011), being a failsafe crop that remains productive in conditions that are too dry for other cereals. Despite its inherent drought tolerance, moisture stress is nonetheless one of the major constraints for sorghum production (Vadez et al., 2011).

Phenotypic selection for drought tolerance is challenging as it is conditioned by complex inheritance and involve a plethora of component traits. A recent study catalogued more than 600 QTL underlying drought and heat stress from ~150 publications in sorghum from 1995 to present (Mace et al., 2019). Transcriptomic analysis comparing stay-green and senescent sorghum genotypes identified more than 2,000 differentially expressed genes, of which 289 were within known stay-green QTL (Johnson, Cummins, Lim, Slabas, & Knight, 2015). Further, plant height and flowering time are extensively studied adaptive traits in sorghum (Bouchet et al., 2017; Brown et al., 2006; Kong et al., 2018; Lin, Schertz, & Paterson, 1995; Morris et al., 2013; Zhang et al., 2015), and along with variation in stay-green and other drought-related traits, are also predictive of grain yield performance in drought prone environments (Jordan et al., 2003).

Here, we explore the potential to build on the inherent drought tolerance of an elite cultivar bred for Ethiopian conditions, by crossing 12 diverse founder lines with different drought adaptations (Vadez et al., 2011) to an Ethiopian elite cultivar, and phenotyping the resulting 12 populations under natural drought environments in Ethiopia, taking advantage of the ‘nested association mapping’ (NAM) strategy (Yu *et al*. 2008) to combine high mapping resolution and high statistical power in studying variation from diverse parental inbreds. Head exsertion provides a valuable indicator of pre-flowering drought stress (Rosenow & Clark, 1981; Seetharama, Reddy, Peacock, & Bidinger, 1982) -- driven by elongation of the uppermost internode of the stem (peduncle) shortly before anthesis, increased peduncle exsertion was also associated with drought resistance in rice (*Oryza sativa* L.) (O’Toole & Namuco, 1983); and was positively correlated with yield, length of head, and number of productive tillers in pearl millet (*Pennisetum americanum* L.) (Ibrahim, Marcarian, & Dobrenz, 1986). The ‘stay-green’ trait, reflecting post-flowering drought tolerance by the ability to maintain green leaf area, resist premature plant senescence, resist lodging, and fill grain normally (Rosenow, Quisenberry, Wendt, & Clark, 1983; Thomas & Ougham, 2014), has been a focus in sorghum (Borrell et al., 2000, 2001, 2014; Crasta et al., 1999; Harris et al., 2007; Kassahun et al., 2010; Sanchez et al., 2002; Subudhi et al., 2000; Xu et al., 2000; Vadez et al., 2011), wheat (*Triticum aestivum*) (Christopher, Manschadi, Hammer, & Borrell, 2008; Lopes & Reynolds, 2012), maize (*Zea mays*) (Campos, Cooper, Habben, Edmeades, & Schussler, 2004; Zheng et al., 2009), and rice (*Oryza sativa*) (Cha et al., 2002; Jiang et al., 2007).

## Materials and Methods

### Plant materials and methods

Twelve donor sorghum lines from eight countries including Nigeria (IS10876), South Africa (IS14298), Sudan (IS14446, IS3583, IS9911), Ethiopia (IS14556), Cameroon (IS15428, IS16044, IS16173), India (IS2205), Botswana (IS22325), and Yemen (IS23988) (Table 1) were obtained from ICRISAT, crossed to the Ethiopian elite cultivar Teshale and the F_1_ plants backcrossed to manually emasculated Teshale to produce BC_1_F_1_ families which were selfed by single seed descent to produce BC_1_F_4_ populations of 32-153 lines. For simplicity, we use the name of the alternate parent to refer to an individual BC_1_F_4_ population (*e*.*g*., the IS10876 population).

**Table 1.** Description of the BC-NAM population

| Name | Country of origin§ | Kobo |  | Meiso |  | Sheraro |  |
| --- | --- | --- | --- | --- | --- | --- | --- |
|  |  | Pop | No. lines | Pop | No. lines | Pop | No. lines |
|  |  | size | with plant loss (%) | size | with plant loss (%) | size | with plant loss (%) |
| IS10876 | Nigeria | 135 | 39 (29%) | 151 | 27 (18%) | 153 | 0 (0%) |
| IS14298 | South Africa | 121 | 48 (40%) | 132 | 16 (12%) | 131 | 1 (1%) |
| IS14446 | Sudan | 134 | 50 (37%) | 145 | 22 (15%) | 149 | 0 (0%) |
| IS14556 | Ethiopia | 32 | 14 (44%) | 34 | 2 (6%) |  |  |
| IS15428 | Cameroon | 124 | 43 (35%) | 142 | 25 (18%) | 144 | 0 (0%) |
| IS16044 | Cameroon | 36 | 9 (25%) | 37 | 2 (5%) |  |  |
| IS16173 | Cameroon | 98 | 25 (26%) | 111 | 18 (16%) | 112 | 0 (0%) |
| IS2205 | India | 36 | 20 (56%) | 40 | 6 (15%) |  |  |
| IS22325 | Botswana | 101 | 33 (33%) | 120 | 25 (21%) | 120 | 0 (0%) |
| IS23988 | Yemen | 34 | 15 (44%) | 42 | 10 (24%) | 36 | 1 (3%) |
| IS3583 | Sudan | 104 | 34 (33%) | 119 | 22 (18%) | 119 | 0 (0%) |
| IS9911 | Sudan | 76 | 17 (22%) | 82 | 14 (17%) | 82 | 0 (0%) |
| Total |  | 1031 | 347 (34%) | 1155 | 189 (16%) | 1046 | 2 (0%) |

□ Founder lines used in the BC-NAM population. Note that population IS32234 was discarded due to severe contamination.
§ Source information is obtained from the International Crops Research Institute for the Semi-Arid Tropics (ICRISAT): http://genebank.icrisat.org/IND/Passport?Crop=Sorghum.

### Experimental design and field managements

Parental lines and the BC-NAM population were evaluated at three locations that represent major sorghum growing regions in Ethiopia (Table 1, Figure S1): Kobo (12°09’N, 39°38’E) and Meiso (09°14’N, 40°45’E) in the 2017 cropping season, and Sheraro (14°23’N, 37°46’E) in the 2018 cropping season. These three locations are typical of the range of moisture stress conditions in the target region and had no irrigation, subjecting the study populations to natural moisture stress conditions. An alpha lattice design with two replications was used at each location. Each field trial was conducted from July to December. Populations IS2205, IS14556, IS16044, IS32234 were not included in Sheraro due to space constraints. Seeds of each sorghum line were sown into a 4 m row and thinned three weeks after sowing to spacings of 75 cm and 20 cm between and within rows, respectively. Therefore, an individual plot without plant loss would consist of 20 plants. During sowing and three weeks after sowing, inorganic fertilizers DAP and Urea were added at the rate of 100 and 50 kg ha^-1^, respectively. Field weeding and other cultural practices were carried out as needed.

### Phenotypic data analysis

A total of eight traits were evaluated. Leaf senescence (LS) was scored on a plot basis at physiological maturity on a 1-5 scale based on degree of leaf death, with 1 = least senescent (10%), 2 = slightly senescent (25%), 3 = intermediate senescent (50%), 4 = mostly senescent (75%), and 5 = completely senescent (100%). Number of leaves (NL) per plant was averaged from five plants per plot. As the lower leaves of sorghum tend to die and fall off as plants mature, all leaves on the main stem were tagged when plants were at five or six leaf stage, so they could be reflected in counts even if they had been shed. Head exsertion (HX) beyond the ligule of the flag leaf was recorded in centimeters. Number of basal tillers (NT) per plant was counted. Thousand seed weight (TSW) was determined from 1,000 randomly selected seeds in grams. Single plant-based grain yield (SGY) was recorded as the total grain weight in grams of threshed panicles of one plant, averaged from five plants per plot. However, due to plant loss in many plots, particularly at Kobo and Meiso fields, individual plant grain yield estimates were confounded by plant density among plots (i.e., plants from low density plots would have received more resources such as light and moisture due to less competition than plants from high density plots). Therefore, counts of plants per plot (CPP) were recorded and used as covariate to adjust for SGY as follows:

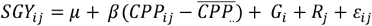

Where *SGY*_*ij*_ represents the original single plant-based grain yield,μ is the grand mean, β is the regression coefficient of the covariate on the genotype, *CPP*_*ij*_ is the value of the covariate for a particular plot, and 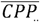 is the overall mean of the covariate, *G*_*i*_ represents genotype, *R*_*j*_ represents replication, and *ε*_*ij*_ is random error. Plot-based grain yield (PGY) was calculated by multiplying adjusted single plant-based grain yield (adjusted SGY) by the number of plants per plot (CPP). Thus, plot-based grain yield would reflect the actual grain yield potential of sorghum genotypes under the targeted natural environments by considering planting density. In Sheraro, LS, HX, and NT did not express symptoms or exhibited limited phenotypic variations and thus were not phenotyped (K. Bantte, personal communications).

Raw data from each location were checked and outlier phenotypes were discarded. HX at Meiso exhibited left-skewed data distribution (Figure S2) and values used in genetic analyses were transformed to approximate normal distribution using Box-Cox method (Box & Cox, 1964), calculated as (*y*^λ^ 1)/λ, where y is the original phenotypic value, and λ is the optimal value which results in the best approximation of normalization. λ was determined as 0.4 in this case. Many genotypes had no exsertion (HX=0), and a constant of one was added because Box-Cox transformation only works if all data are positive. To interpret results such as allelic effects, transformed data were converted back to the original values.

In order to evaluate the effects of environment, family, genotype nested in family, family by environment interaction, and genotype nested in family by environment interaction, a multi-environment analysis of variance (ANOVA) was conducted for each trait across all three environments. Due to the strong environment and genotype-by-environment effects (Table S1), trait best linear unbiased predictions (BLUPs) were estimated for each line within each environment using a mixed linear model implemented in the lme4 package (Bates, Mächler, Bolker, & Walker, 2015). All model terms were treated as random effects in the following model:

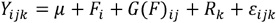

where *Y* represents raw phenotypic data, *F* is the individual family in BC-NAM population,*G*(*F*) is genotype nested within family, *R* is replication, and *ε* is random error. Pearson correlation coefficients between traits were calculated using line BLUPs. Broad-sense heritability in each environment was determined as the proportion of total phenotypic variance explained by the combined family and line terms using equation 1 from Wallace *et al*. (2016):

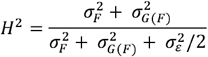

where 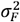 the variance explained by family, 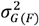 is the variance explained by individual lines, and 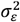 is the random error variance.

### Marker-trait association

A total of 4395 SNPs generated using a *Pst*I-*Msp*I protocol were used for marker-trait association analyses. Details of marker development and filtering were described as follows: GBS libraries were sequenced on a NextSeq500 (Illumina, San Diego, CA) with 150-bp single-end reads at the Genomics and Bioinformatics Core of the University of Georgia; SNP calling was conducted using TASSEL GBS v2 pipeline (Bradbury et al., 2007) and *Sorghum bicolor* genome v2.1 (Paterson et al., 2009); then raw SNPs were filtered based on tag coverage > 5%, minor allele frequency > 0.03, single marker missing data < 5%, yielding a set of 4395 SNPs for analysis. QTL mapping considered both joint-linkage (JL) (Buckler et al. 2009) and GWAS models. JL analysis only included eight large populations (N > 80, Table 1) by removing IS14556, IS16044, IS2205, and IS23988, which contain many monomorphic SNPs and pose a problem in accurately estimate marker effects due to their small size. JL mapping used the forward-backward stepwise regression model implemented in the Stepwise Plugin of TASSEL 5, with family as a cofactor and marker nested within family, suggested to be the most powerful procedure for JL mapping (Würschum et al., 2012). The family main effect was fit first, then marker effects were selected to enter or leave the model based on the E-03 *P*-value threshold for the marginal *F*-test of that term. GWAS was performed using the lm function in R (R Development Core Team 2016) by fitting SNPs as non-nested effects with family as a cofactor. All 12 populations were included in GWAS because SNPs were non-nested and treated as biallelic, thus small size of individual populations was less a problem. Loci of the GWAS model that had *P*-value below the E-03 threshold were considered QTL.

All JL QTLs identified in this study were compared to the Sorghum QTL Atlas database [Mace *et al*. (2019)], which collated the projected locations of *c*. 6000 QTL or GWAS loci from 150 publications in sorghum from 1995 to present. Peak SNP associations from JL and GWAS were also compared to *c*. 2000 genes differentially expressed between stay-green (B35) and senescent (R16) sorghum genotypes (Johnson et al., 2015). Sorghum genes containing or directly adjacent to SNP associations were also searched using BEDOPS (Neph et al., 2012) and standard UNIX tools.

### Genome-wide prediction

Drought-related traits are predictive of grain yield in drought prone sorghum production environments (Jordan et al., 2003). We performed genomic predictions for plot-based grain yield (PGY) and 1000-seed weight (TSW) using marker-trait associations for the eight drought responsive traits in this study and three adaptive traits mapped in a companion study of this sorghum BC-NAM population including days to flowering, days to maturity, and plant height (Dong et al., unpublished data). Specifically, we considered the following three scenarios for genomic prediction: (i) All 4395 segregating SNPs across the genome; (ii) the peak SNPs identified from JL and GWAS of all eleven traits from three environments; (iii) an equal number of random SNPs that have the similar allele frequencies as those detected SNPs from JL and GWAS. All models were run with 50 five-fold cross validations, in which 20% of phenotypes were randomly masked. The resulting predictions were compared to the masked (true) values to generate a prediction accuracy estimate, measured using a Pearson correlation coefficient. *P*-values for the differences in distribution of correlations were determined via a two-sided *t*-test.

## Results

### Phenotypic variation within and between families

Different natural drought conditions provided opportunities to study this BC-NAM population across three environments. Multi-environment ANOVAs revealed strong and significant effects of environment, family, genotype nested in family, family by environment interaction, and genotype by environment interaction for all traits, expect that genotype by environment effect was not significant for head exsertion (Table S1). During the 2017 cropping season, Kobo and Meiso experienced severe moisture stress during sowing season (July). Daily data from nearby weather stations showed cumulative precipitation before sowing (Jan-June) of 194.4 mm and 313.6 mm in Kobo and Meiso, respectively, and 401.2 mm in Sheraro (Figure S1). At Kobo, a total of 347 lines (34%) across all 12 families had plant loss, ranging from 22% in IS9911 to 56% in IS2205 (Table 1). At Meiso, 189 lines (16%) lost one of the two replicates, and plant loss percentage ranged from 5% in IS16044 to 24% in IS23988. Between plant loss at Kobo and Meiso there were 58 lines in common, however neither the Pearson’s correlation coefficient (*r* = 0.021, *p* = 0.9484) nor the rank-based Kendall’s correlation statistic (τ = 0.030, *p* = 0.9466) were significant, indicating that sorghum response to drought is dynamic and labile under different stressed environments. However, during the 2018 cropping season in Sheraro, seeds germinated well and virtually all plants survived (Table 1).

Phenotypic distribution and variance differed greatly among the three environments (Figure 1). Five traits including leaf number, single plant-based grain yield, count of plants per plot, plot-based grain yield, and 1000-seed weight were evaluated over three locations; three traits including head exsertion, leaf senescence, and tiller number were evaluated at Kobo and Meiso only (Figure 1). Plants grew fewer leaves in Kobo (avg. leaf number: 8.76) and Meiso (avg. leaf number: 9.95) than Sheraro (avg. leaf number: 13.60) (Table S2, Figure 1). Head exsertion and leaf senescence are two interesting traits to evaluate drought tolerance in sorghum, the former trait showed larger phenotypic variation in Meiso, whereas the latter trait exhibited more phenotypic variations in Kobo (Figure 1), presumably reflecting somewhat different drought regimes. Phenotypic variation of tiller number was quite limited in this BC-NAM population, especially in Meiso (Figure 1).

**Figure 1.**
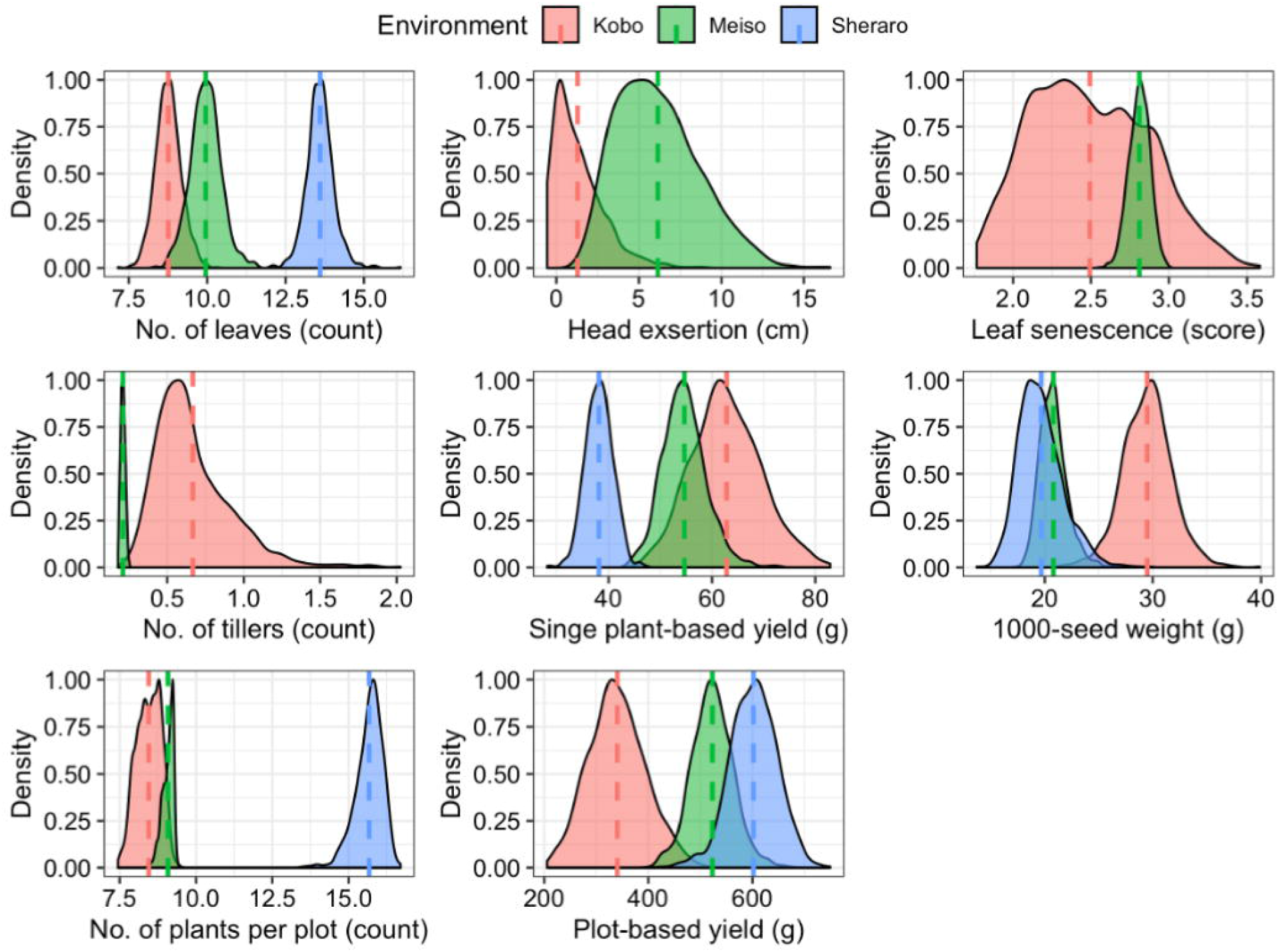
Phenotypic distribution of eight traits in the BC-NAM population at three environments. x axis represents trait metrics and y axis represent distribution density. Vertical dashed lines represent population means at respective environment. NL: number of leaves, HE: head exsertion, LS: leaf senescence, NT: number of tillers, SGY: single plant-based grain yield, CPP: count of plants per plot, PGY: plot-based grain yield, TSW: 1000-seed weight.

**Figure 2.**
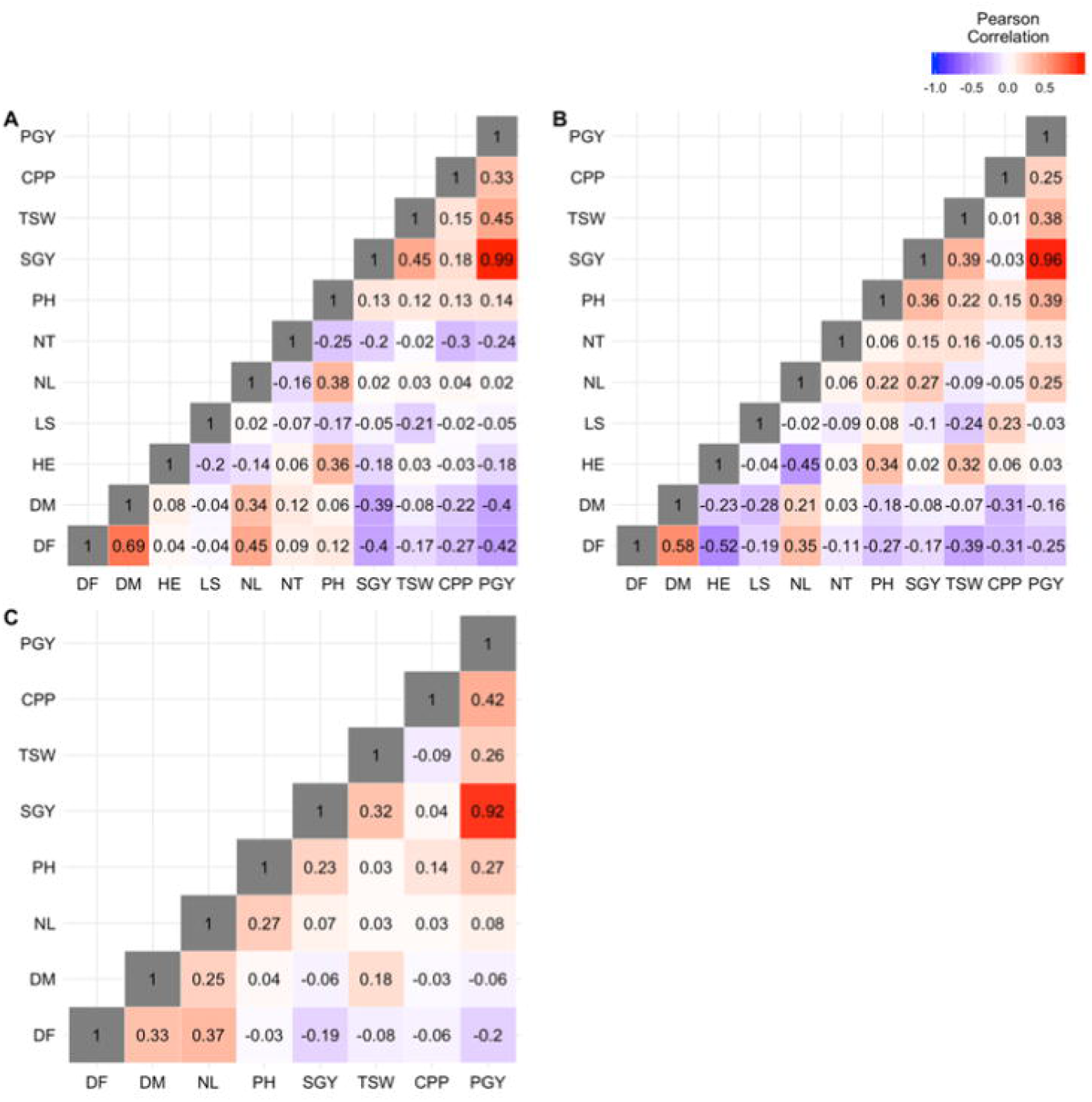
Trait correlations among nine traits studied in this BC-NAM population. Note that in addition to the eight traits in this study, we also included three other traits including days to flowering (DF), days to maturity (DM), and plant height (PH), that were genetically dissected in a parallel study.

Plot-based grain yield was highest in Sheraro, followed by Meiso, and lowest in Kobo (Figure 1), reflecting relative levels of plant loss and plant density at maturity. In contrast, single plant-based grain yield and 1000-seed weight ranked highest in Kobo, followed by Meiso, and lowest in Sheraro (Figure 1, Table S2). The nonuniform plant density across three environments might also lead to the differences of length of grain filling period (anthesis to maturity) at three locations -- subtracting days to flowering from days to maturity, plants in Kobo (48.89 d.) and Meiso (37.61 d) had significantly longer grain filling periods than in Sheraro (25.23 d; Table S2).

While many phenotypes of individual families were similar to that of the recurrent parent, Teshale (Figure S3, File S1), consistent with the backcross breeding scheme, all families showed higher head exsertion and lower leaf senescence indicating improved pre- and post-flowering drought tolerance in the most stressed tests. In Kobo, population means for head exsertion ranged from 0.57 cm in IS2205 to 2.3 cm in IS22325 (Figure S3, File S1), whereas that of recurrent parent Teshale was only 0.03 cm; in Meiso, population means for head exsertion ranged from 4.95 cm for IS9911 to 7.60 cm in IS15428, whereas Teshale was 3.92 cm (Figure S3, File S1).

All 12 families also showed lower leaf senescence mean values than the recurrent parent at Kobo and Meiso (Figure S3), based on an ordinal scale 1-5 to evaluate leaf senescence with 1 being least senescent and 5 being complete senescent. In Kobo, population means for leaf senescence ranged from 2.30 in IS14298 to 2.64 in IS14556, whereas that of Teshale was 2.67; in Meiso, population means for leaf senescence ranged from 2.79 cm for IS2205 and IS9911 to 2.84 in IS14556 and IS16044, whereas that of Teshale was 2.86 (File S1). Therefore, phenotype data of head exsertion and leaf senescence indicated that drought tolerance alleles from diverse founder lines could have been introgressed into the recurrent parent.

Broad-sense heritabilities of the eight traits across the BC-NAM population within each environment varied from 0.08 for leaf senescence in Meiso to 0.63 for 1000-seed weight in Sheraro (Table 2). Head exsertion showed moderate heritabilities at Kobo (0.58) and Meiso (0.59). Heritability of leaf senescence was moderate in Kobo (0.42) but very low in Meiso (0.28), which was primarily due to the limited variation in Meiso (Figure 1). Number of tillers was similarly low heritable at both Kobo (0.27) and Meiso (0.26). Number of leaves showed moderate heritabilities across three environments (0.44-0.60). Plot-based grain yield was less heritable than 1000-seed weight, and heritabilities of these two traits ranged from 0.26-0.39 and 0.31-0.63, respectively. Count of plants per plot, which was used as the covariate to adjust for planting density in the estimation of plot-based grain yield, exhibited low heritabilities across three environments (0.16-0.38).

**Table 2.** Summary of joint-linkage QTL of each trait in the BC-NAM population

| Trait§ | Kobo |  |  | Meiso |  |  | Sheraro |  |  |
| --- | --- | --- | --- | --- | --- | --- | --- | --- | --- |
| | $H^2$¶ | No. QTL | Model $R^2$□ | $H^2$ | No. QTL | Model $R^2$ | $H^2$ | No. QTL | Model $R^2$ |
| HE | 0.58 | 7 | 0.26 (45%) | 0.59 | 12 | 0.37 (63%) |  |  |  |
| LS | 0.42 | 3 | 0.21 (50%) | 0.28 | 9 | 0.21 (75%) |  |  |  |
| NT | 0.27 | 7 | 0.19 (70%) | 0.26 | 10 | 0.24 (92%) |  |  |  |
| NL | 0.44 | 13 | 0.33 (75%) | 0.49 | 6 | 0.18 (37%) | 0.60 | 8 | 0.19 (32%) |
| SGY | 0.34 | 4 | 0.12 (35%) | 0.28 | 8 | 0.19 (68%) | 0.28 | 4 | 0.11 (39%) |
| CPP | 0.38 | 2 | 0.10 (26%) | 0.16 | 5 | 0.14 (88%) | 0.18 | 2 | 0.06 (33%) |
| PGY | 0.39 | 10 | 0.35 (90%) | 0.26 | 8 | 0.23 (88%) | 0.28 | 6 | 0.18 (64%) |
| TSW | 0.59 | 6 | 0.18 (31%) | 0.31 | 10 | 0.24 (77%) | 0.63 | 14 | 0.31 (49%) |

§ HE: head exsertion; LS: leaf senescence; NL: number of leaves; NT: number of tillers; SGY: single plant-based grain yield; CPP: count of plants per plot; PGY: plot-based grain yield; TSW: thousand seed weight.
¶Broad-sense heritability of BC-NAM population.
□Variance explained by joint-linkage model after fitting family term and detected QTL.
Numbers in parentheses represent proportion of genetic variance explained by the join-linkage model, which was calculated by diving model $R^2$ by $H^2$ .

### Genetic architecture of adaptive traits

Genetic architecture of complex traits is defined by the number, effect size, frequency, and gene action of the QTLs that affect it. We conducted association analyses for eight adaptation traits. A total of 154 JL QTLs were detected using joint-linkage model across the three environments (Table 2, Figure 3A, File S2), of which 100 (64.9%) overlapped with previous QTLs detected for the same trait from 72 different studies (File S2). Large number of common QTLs of same traits between this study and many previous studies demonstrated the power of this sorghum BC-NAM population in dissecting drought tolerance traits by sampling more allelic diversity than possibly within any single biparental population. In order to leverage the investment, we incorporated the four small populations (IS2205, IS14556, IS16044, and IS32234) into GWAS analyses. A total of 233 GWAS associations were detected, with 103, 90, and 40 in Kobo, Meiso, and Sheraro, respectively (Figure 3A, File S3). Despite the different statistical frameworks, high correspondence between JL QTLs and GWAS hits was observed (Figure 3A). Exact overlap of linkage mapping and GWAS is not expected, as linkage mapping tests markers within an individual population whereas GWAS tests marker effects across populations, leading to different strengths and weakness in each approach (Tian et al., 2011). Joint linkage analysis produces many more small effects than GWAS analysis as an artifact of the model fitting process, which assigns a separate effect to all populations at each QTL. Moreover, the addition of those four small populations also played a role in the discrepancy of associations between these two methods.

**Figure 3.**
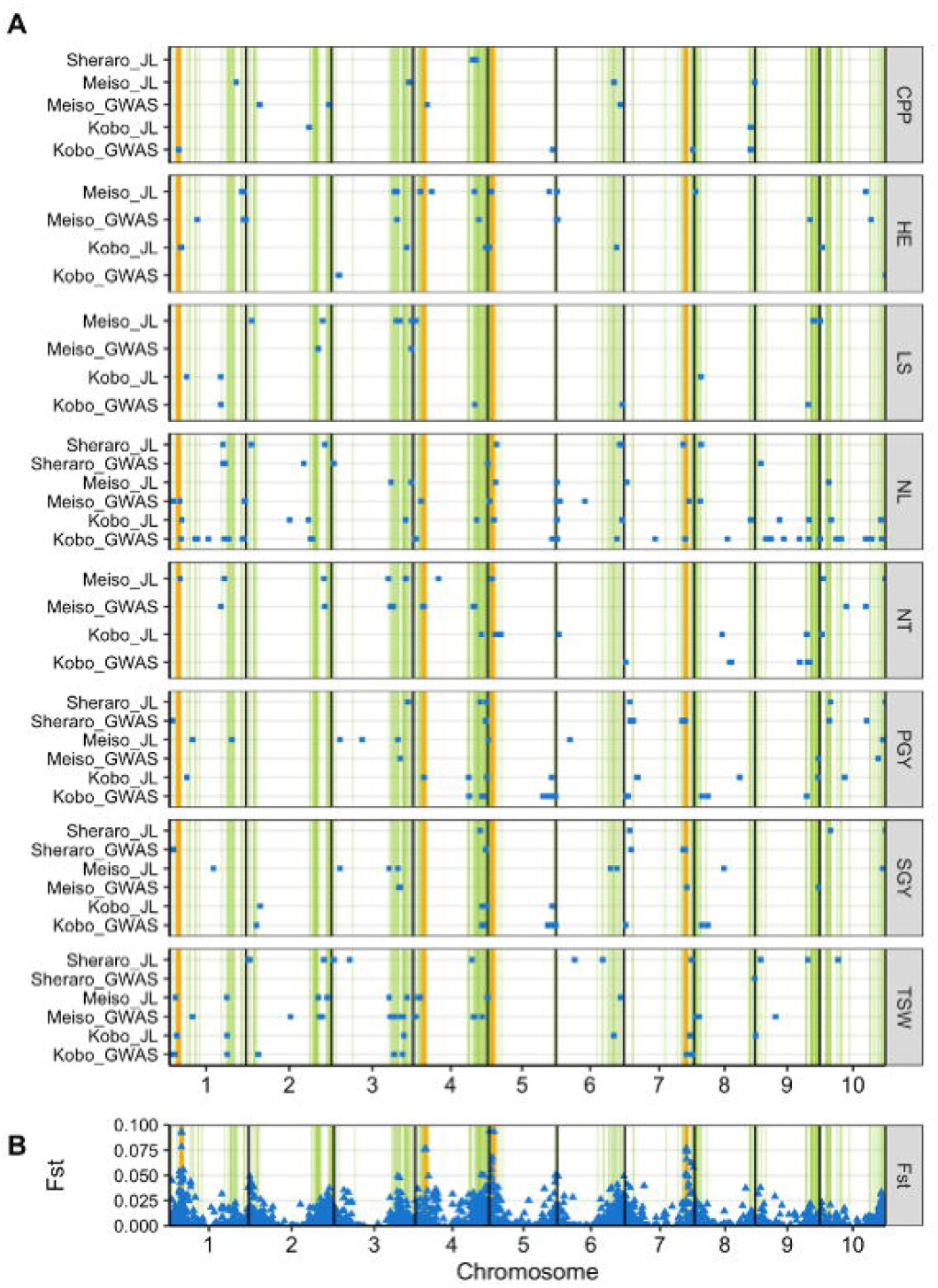
Genomic regions associated with drought-related traits in the sorghum BC-NAM population. (A) Distribution of the 154 joint-linkage QTLs and 233 GWAS hits for six traits, which were represented with blue dots. NL: number of leaves, HE: head exsertion, LS: leaf senescence, NT: number of tillers, SGY: single plant-based grain yield, CPP: count of plants per plot, PGY: plot-based grain yield, TSW: 1000-seed weight. (B) Fixation index (*F*_*ST*_) between the 58 drought-sensitive lines, which lost one of two replicates at both Kobo and Meiso, and the remaining 1113 lines. Orange vertical lines in both panels represent *F*_*ST*_ peaks.

Genes within stay-green QTLs were enriched for proximity to drought-related trait loci identified in this study (Figure 3). The 154 JL QTLs and 233 GWAS hits were distributed across all ten chromosomes in a non-random fashion (Figure 3A). We compared these associations with the transcriptomic analysis comparing stay-green and senescent sorghum lines (Johnson *et al*. 2015), which detected 2,036 differentially expressed genes between the stay-green (B35) and senescent (R16) sorghum genotypes. In particular, the associations detected in this study clustered around 289 genes within known stay-green QTLs catalogued in the Comparative Saccharinae Genome Resource (File S4) (Zhang et al., 2013), which further corroborated the reliability of the associations detected in this study. Due to the moisture stress during sowing season (Figure S1), 58 sorghum lines lost one of their two replicates at both Kobo and Meiso (Table 1), indicating that these lines may be more sensitive to moisture stress than the remaining 1113 lines. Intriguingly, fixation index (*F*_*ST*_) analysis revealed elevated peaks at stay-green QTL regions between these 58 drought-sensitive lines and the other 1113 lines (Figure 3B), on chromosome 1 (*F*_*ST*_ = 0.093, S01_11158453), chromosome 4 (*F*_*ST*_ = 0.077, S04_9538999), chromosome 5 (*F*_*ST*_ = 0.094, S05_363813), and chromosome 7 (*F*_*ST*_ = 0.077, S07_56408236). Many associations from joint-linkage and GWAS analyses were also detected near these *F*_*ST*_ peaks (Figure 3A).

Allelic effect of QTLs is an important facet of genetic architecture of traits. For the 154 JL QTLs, marker effects within each individual population were estimated (i.e. 154 QTLs × 8 populations = 1,232 QTL effects). Since most QTLs were not present in all families, many of these effects were near zero and generally normally distributed (Figure S4, File S2). Fisher (1930) provided a simple theoretical justification for these observations. For a well-adapted organism close to its fitness optimum, only small effects can increase fitness. This explanation conforms to the breeding design of this BC-NAM population, that all founder lines and the common recurrent parent are generally adapted to semiarid environments. In addition, Orr (1998) showed that regardless of the distance from the fitness optimum, the expected distribution of effect sizes progressively fixed during adaptation is exponential, with a small number of large-effect loci fixed first, followed by progressively larger numbers of loci with small effects becoming fixed. Moreover, the genetic architecture of intraspecific variation like BC-NAM population in this study consists of many loci with small effects because loci with larger effects tend to be only briefly polymorphic.

Among the seven head exertion JL QTLs at Kobo, allelic effects ranged from −2.9 cm at S01_14044648 in IS22325 to 2.8 cm at S01_12672806 in IS22325 (Figure S4, File S2); 34 showed positive effects (i.e., increase head exertion ability) versus 22 negative allelic effects. Among the 12 JL QTLs for head exertion in Meiso, allelic effects ranged from −1.3 cm at S05_55829561 in IS14446 to 1.1 cm at S01_71318983 in IS22325 (Figure S4, File S2). Leaf senescence allelic effects were of small magnitude, ranging from −0.6 (i.e., negative number means less senescent) at S01_49654172 in IS3583 to 0.5 (i.e., positive number means more senescent) at S08_5990097 in IS10876 at Kobo; allelic effects of leaf senescence ranged from −0.11 at S09_54402475 in IS9911 to 0.08 at S10_439084 in IS16173 in Meiso (File S2). Smaller allelic effects of leaf senescence in Meiso compared to those in Kobo corresponds to the narrower trait phenotypic variation in Meiso (Figure 1). For number of leaves, allelic effects ranged from −1.3 (i.e., negative number means less leaves than recurrent parent) at S08_51534279 in IS22325 to 0.8 at S10_10472812 in IS22325 at Kobo; and allelic effects ranged from −0.7 at S07_2397109 in IS9911 to 0.8 at S06_769807 in IS14298. For number of tillers, allelic effects ranged from −0.3 (i.e., negative number means reduced tillering compared to recurrent parent alleles) at S04_62057354 in IS13428 to 0.3 at S05_11960542 in IS10876 at Kobo; whereas in Meiso, allelic effects for number of tillers were very small, ranging from −0.1 to 0.2, which again corresponds to the very limited trait phenotypic variation in Meiso (Figure 1). Such approximately normally distributed allelic effects were observed for all traits studied (Figure S4). Indeed, there were QTLs exhibiting large allelic effects within specific families (Figure S4). For example, at one plot-based grain yield QTL detected in Kobo on chromosome 4 (S04_50497689), the IS9911 allele contributed a positive effect of 97.44 g relative to the recurrent parent (File S2), which is considerably very large given that plot-based grain yield across the whole BC-NAM population ranged from 205.81 g to 512.03 g in Kobo (File S1). Large effect alleles in this study were of great breeding value in improving Ethiopian adapted cultivars.

Total phenotypic variance explained by the final JL model was generally low for these eight adaptive traits, ranging from 0.10 for CPP to 0.35 for PGY in Kobo, from 0.14 for CPP to 0.37 for HE in Meiso, and from 0.06 for CPP to 0.31 for TSW in Sheraro (Table 2). Note that heritability imposes an approximate upper limit to the *R*^2^ of a QTL model (Yu et al., 2008). Therefore, this is not unexpected given the low to moderate heritabilities of these traits under drought conditions. By taking the broad-sense heritability into account, proportion of genetic variance explained by the final joint-linkage model of each trait ranged from 26% for CPP to 90% for PGY in Kobo, 37% for NL to 88% for LS and PGY in Meiso, and 32% for NL to 64% for PGY in Sheraro (Table 2).

Some of the strongest and (or) most consistent associations of various traits locate near known genes. Green leaf is the hub of photosynthesis and leaf number has been of particular interest in drought tolerance studies in sorghum. JL model detected 13, 6, and 8 QTLs for leaf number in Kobo, Meiso, and Sheraro, respectively (Table 2, Figures 3, 4). Significant associations for leaf number were located near the *Ma6* region at Kobo and Meiso, with QTL peaks (S06_782670 at Kobo and S06_769807 at Meiso) 109 kb and 96 kb away from *Ma6*, respectively (Figure 4A and 4B). However, this association was not detected in Sheraro (File S2). In addition to flowering regulators, associations were also detected near known stress regulation genes. One *DREB1A* gene (Sobic.007G181500), which encodes a dehydration-responsive element-binding transcription factor, was located at the *F*_*ST*_ peak region on chromosome 7 (*F*_*ST*_ = 0.077, S07_56408236), and GWAS association for leaf number at Meiso (S07_59324541; Figure 4B, File S3), and JL-QTL for 1000-seed weight at Kobo (S07_60171225; File S2) were detected near this gene. Several traits including number of leaves (JL-QTL S03_67359281 at Kobo, Figure 4A, File S2), head exsertion at Meiso (S03_59654236, Figure 4C, File S2), and number of tillers at Meiso (S03_67407460 at Meiso, File S2) also showed strong associations near the *P5CS2* gene (delta1-pyrrolinr-5-carboxylate synthase 2; Sobic.003G356000, File S2). Among these, the association for number of tillers in Meiso (S03_67407460) was only 33 kb from *P5CS2*, encoding an enzyme responsible for proline biosynthesis (Kishor, Hong Zonglie, Miao Guo-Hua, Hu Chein-An, & Verma, 1995) that is highly expressed in the stay-green sorghum line compared with the senescent line (Johnson et al., 2015). Several traits including head exsertion, number of leaves, number of tillers, and 1000-seed weight had associations on chromosome 5 (Figure 4, File S2, File S3) near the *LEA* gene (Sobic.005G021800) encoding hydrophilic proteins with major roles in drought and other abiotic stresses tolerance in plants (Magwanga et al., 2018). The *LEA* gene on chromosome 5 was also located near the *F*_*ST*_ peak (*F*_*ST*_ = 0.094, S05_363813, Figure 3) distinguishing the 58 sorghum lines lost one of their two replicates at both Kobo and Meiso from the remaining 1113 lines.

**Figure 4.**
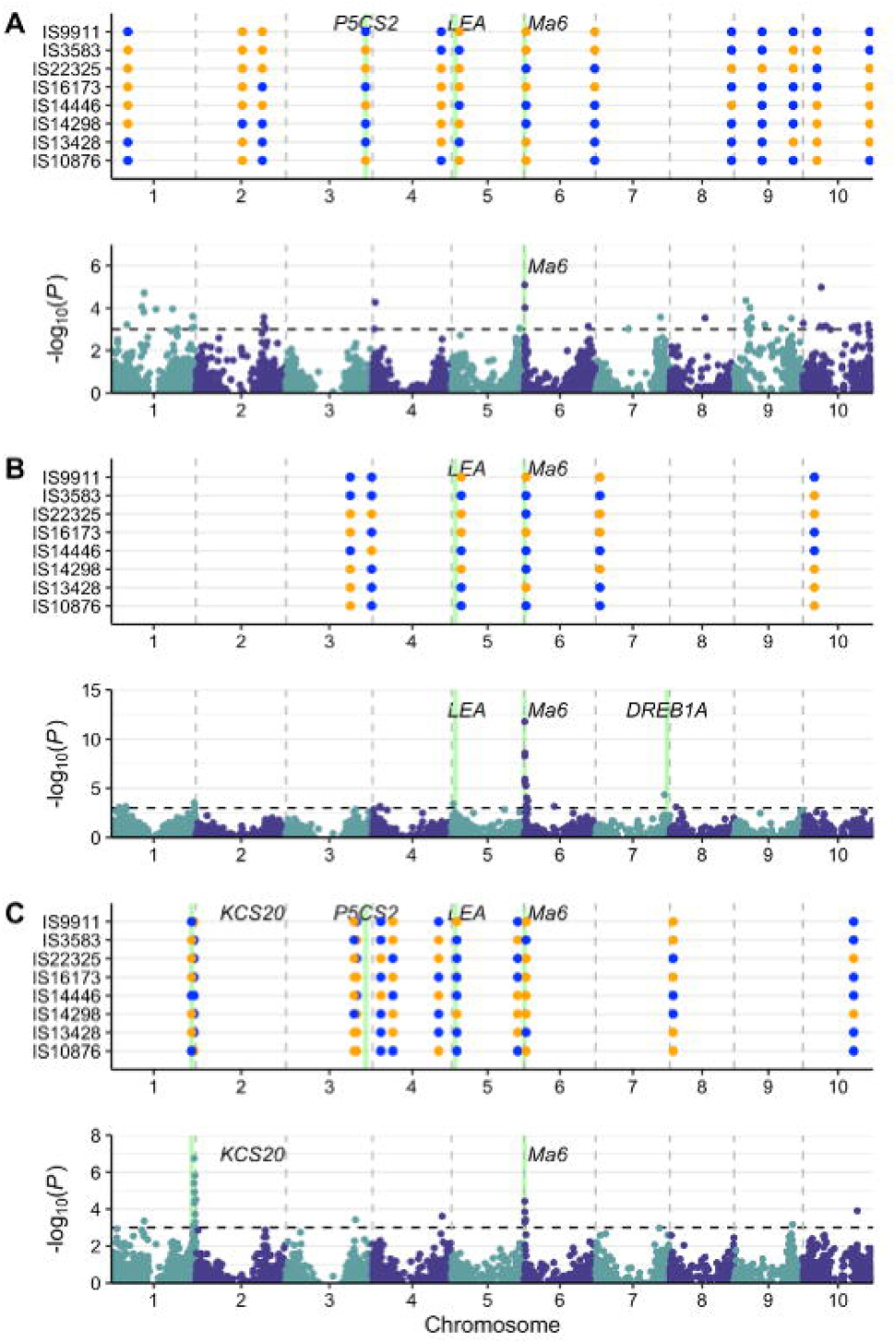
Marker-trait associations of drought-related traits. Each panel shows the associations detected in joint-linkage model (top) and in GWAS model (bottom). (A) number of leaves in Kobo, (B) number of leaves in Meiso, and (C) head exsertion in Meiso. In joint-linkage analysis, parental allelic effects were color coded, with blue represents positive effect and orange represents negative effect. Candidate genes were shown in green vertical lines and annotated with respective gene names.

Moreover, significant association for head exsertion in Meiso (S01_69085453) was 217 kb from the *KCS20* gene (3-ketoacyl-CoA synthase 20; Sobic.001G495500, Figure 4C), found to be up-regulated by drought and osmotic stress (Joubès et al., 2008). In *Arabidopsis*, expression of *KCS20* was up-regulated twofold by drought (Lee et al., 2009).

## Discussion

Drought-related traits are very complex and polygenic as evidenced by our association analyses. In genetic dissection of yield and drought-related traits under drought conditions, a major and universally recognized obstacle is properly phenotyping in a high-throughput fashion (Tuberosa, 2012). Phenomics-enabled breeding using high-throughput phenotyping (HTP) may enable researchers to better understand the complexities of trait development and to better optimize genotypes through selection in breeding programs (Jiang et al. 2018; Pugh et al. 2018; Xu et al. 2018).

Most of the associations detected for these eight traits located within known stay-green QTL regions (Figure 3). In sorghum, four major stay-green quantitative trait loci (QTL) have been consistently identified (Crasta et al., 1999; Subudhi et al., 2000; Xu et al., 2000; Sanchez et al., 2002; Harris et al., 2007), with a number of physiological studies testing their effects across different genetic backgrounds (Borrell et al., 2000, 2001, 2014; Kassahun et al., 2010; Vadez et al., 2011). Nearby candidates for these associations could be of great interest for future functional studies. Limited knowledge of drought resilience restricts the power of candidate approaches, but it is nonetheless notable that several previously known drought tolerance genes were associated with drought resilience in this study.

While it is clear that the drought resilience of an elite Ethiopian cultivar can be improved by incorporation of different adaptations from diverse germplasm, none of the 12 individual populations produced higher population mean yield that the recurrent parent Teshale (Figure S3, File S1). The estimated plot-based grain yield of Teshale were 401.05 g and 539.42 g at Kobo and Meiso, respectively; while population mean yield ranged from 309.99 g in IS22325 to 377.63 g in IS16044 at Kobo, and from 506.41 g in IS22325 to 536.99 g in IS16173 at Meiso (File S1). Since the parents were not tested at Sheraro, similar plot yield comparison could not be conducted. Transgressive segregants found in each population (Figure S3, File S1) suggest that the adaptive phenotype of Teshale can be recapitulated or perhaps even further improved, strengthening drought resilience of cultivars that stand between vulnerable geographies and famine. For all traits, rich variation was reflected by mixtures of ‘positive’ and ‘negative’ QTL alleles (Figure S4, File S2), suggesting the 12 diverse founder lines with different drought adaptations (Vadez et al., 2011) complementing and supplementing the inherent drought resilience of Teshale. Correlations of plot-based grain yield across all three environments with days to flowering (−0.20 to −0.42); and plant height (0.14-0.39) exemplify scope for the sorts of adjustments that may be needed to re-establish locally adaptive phenotypes. Indeed, with the enormous altitudinal variation of a country such as Ethiopia, somewhat different lines may be needed for different locales.

Genomic prediction for plot-based grain yield and 1000-seed weight based on our findings support the assertion that drought-related traits are predictive of grain yield performance in drought prone sorghum production environments (Jordan et al., 2003). We conducted genomic prediction across the entire BC-NAM population within each environment under three scenarios: using all 4395 genome-wide markers, using 434 model-selected SNPs from JL and GWAS for 11 traits (eight in this study and three in a companion study including days to flowering, days to maturity and plant height; File S2, File S3), and using an equivalent number (434) of randomly selected SNPs that are at least 260 kb based on LD decay away from JL QTL or GWAS hits. Prediction accuracy was low to modest, with average correlations of 0.16-0.34 for plot-based grain yield and 0.25-0.41 for 1000-seed weight, but the selected 434 SNPs had similar accuracy as all 4395 genome-wide SNPs and were significantly more accurate than 434 random SNPs (Figure 5, File S5). Low to moderate genomic prediction accuracies were not unexpected -- the theoretical upper limit of genomic prediction accuracy cannot exceed narrow-sense heritability because rrBLUP uses the additive kinship matrix among genotypes to perform genomic prediction. In this study, most adaptive traits showed low to moderate heritabilities especially under drought conditions (Table 2).

**Figure 5.**
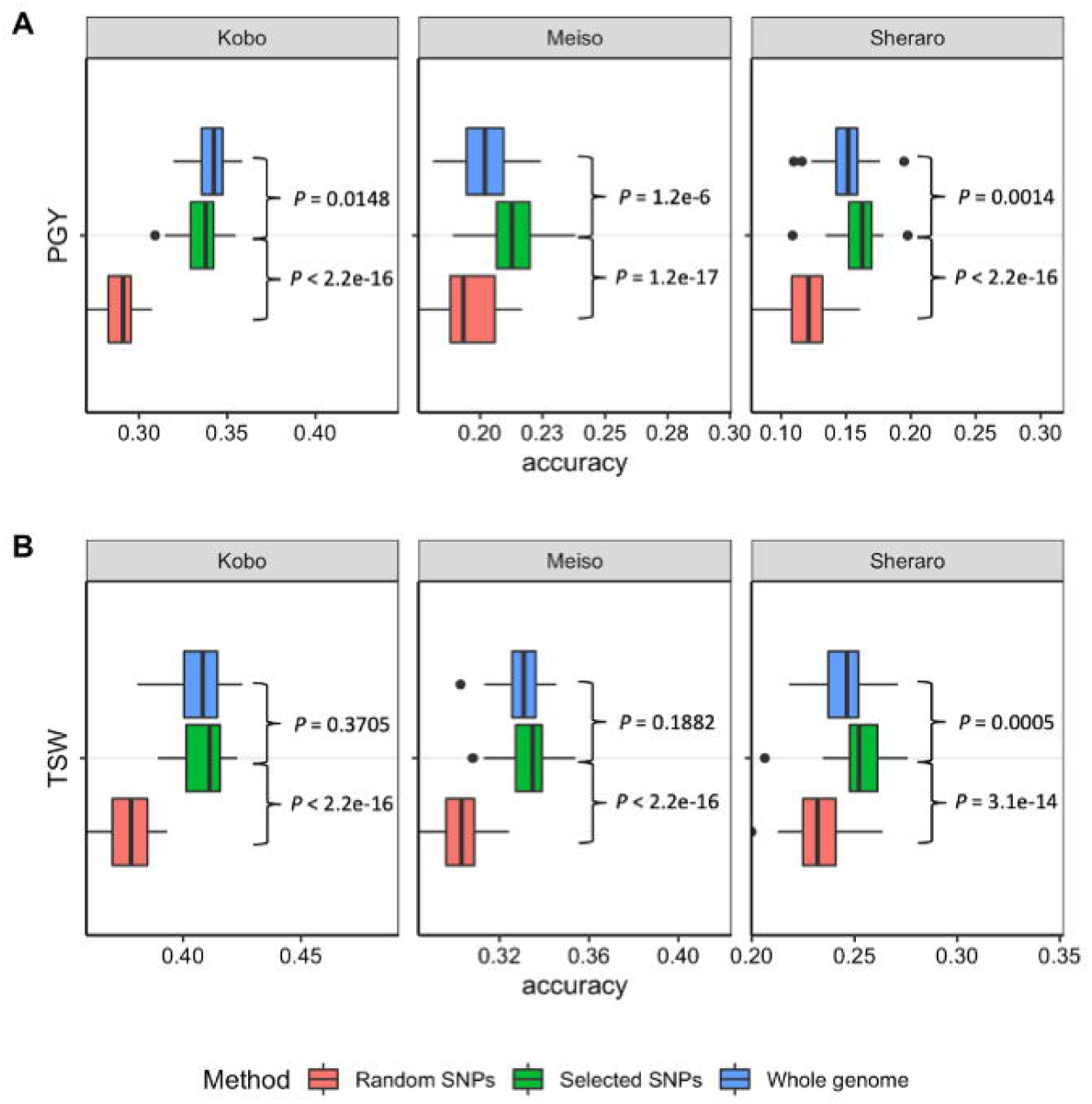
Genome-wide prediction of sorghum grain yield. Phenotypic prediction was performed across the entire sorghum BC-NAM population in 50 five-fold cross-validation using either all 4395 single nucleotide polymorphisms (SNPs), the model-selected SNPs, or an equivalent number of random SNPs. (A) Distribution of genomic prediction accuracy for plot-based grain yield (PGY) across the entire BC-NAM population at three environments. (B) Distribution of genomic prediction accuracy for 1000-seed weight (TSW) across the entire BC-NAM population at three environments. *P* values comparing distributions via a two-sided *t*-test were shown in each plot.

With a worldwide water crisis looming (Rosegrant, Cai, & Cline, 2002; Serageldin, 2004; UNESCO, 2002), increasing the inherent drought resilience of sorghum is vital in rain-fed environments, exemplified by but not limited to sub-Saharan Africa. The sorghum BC-NAM population reported here demonstrates proof of principle, as well as a foundation for breeding improved cultivars for East Africa and a valuable community resource for understanding drought-related traits. Clustering of associations within stay-green QTL regions highlighted the merit of more in-depth exploration of these genomic regions. Given the complex and labile nature of drought-related traits, breeding crops with improved yield and resilience to environmental stresses may greatly benefit from the integration of phenomics technologies such as HTP with genomics.

## Acknowledgments

This work was funded in whole or part by the United States Agency for International Development (USAID) Bureau for Resilience and Food Security under Agreement # AID-OAA-A-13-00044 as part of Feed the Future Innovation Lab for Climate Resilient Sorghum. Any opinions, findings, conclusions, or recommendations expressed here are those of the authors alone. The authors thank Jimma University, Ethiopian Institute of Agricultural Research, Amhara Region Agricultural Research Institute and Tigray Region Agricultural Research Institute for providing land and technical support for the field trials, and the Paterson Lab for valuable help and discussions. We also thank the Georgia Genomics and Bioinformatics Core for sequencing service.

## Data availability statement

Sequencing data are available in the NCBI Sequence Read Archive under project accession BioProject ID PRJNA687679. Data analysis scripts have been deposited to GitHub (https://github.com/hxdong-genetics/Ethiopian-Sorghum-BC-NAM). Please contact corresponding author for other data.

## Figure Legends

**Figure S1** Geographical information and natural environmental conditions of three field trial locations in Ethiopia.

**Figure S2** Geographical information of three field trial locations in Ethiopia

**Figure S3** Phenotypic distribution of eight traits within each BC-NAM family at three environments. x axis represents family names and y axis represent trait metric. Horizontal dashed lines represent recurrent parent means at respective environment.

**Figure S4** Joint-linkage QTL allelic effects distribution for six traits. NL: number of leaves, HE: head exsertion, LS: leaf senescence, NT: number of tillers, SGY: single plant-based grain yield, CPP: count of plants per plot, PGY: plot-based grain yield, TSW: 1000-seed weight.

