## Supplementary material for "Natural variation further increases resilience of sorghum bred for chronically drought-prone environments": Table S1, Table S2

**Table S1.** Analysis of variance of six traits in the sorghum BCNAM population across three environments (Kobo, Meiso, and Sheraro). Phenotypic traits include number of leaves (NL), head exsertion (HE), leaf senescence (LS), number of tillers (NT), single plant-based grain yield (SGY), plot-based grain yield (PGY), and 1000-seed weight (TSW).

| Trait | Term | DF | SS | MS | F | *P* value |
| --- | --- | --- | --- | --- | --- | --- |
| LS | Environment | 1 | 114.4 | 114.36 | 117.418 | < 2e-16 |
|  | Family | 11 | 77.5 | 7.05 | 7.235 | 3.70E-12 |
|  | Genotype | 1147 | 1401 | 1.22 | 1.254 | 1.38E-05 |
|  | Family*Environment | 11 | 64.8 | 5.89 | 6.047 | 9.12E-10 |
|  | Genotype*Environment | 1015 | 1256.4 | 1.24 | 1.271 | 8.95E-06 |
|  | Residuals | 1650 | 1607 | 0.97 |  |  |
| NL | Environment | 2 | 24694 | 12347 | 15591.13 | < 2e-16 |
|  | Family | 11 | 146 | 13 | 16.717 | < 2e-16 |
|  | Genotype | 1159 | 2029 | 2 | 2.211 | < 2e-16 |
|  | Family*Environment | 19 | 53 | 3 | 3.538 | 3.35E-07 |
|  | Genotype*Environment | 2040 | 2616 | 1 | 1.619 | < 2e-16 |
|  | Residuals | 2693 | 2133 | 1 |  |  |
| HE | Environment | 1 | 10981 | 10981 | 616.792 | < 2e-16 |
|  | Family | 11 | 6424 | 584 | 32.8 | < 2e-16 |
|  | Genotype | 1147 | 42996 | 37 | 2.105 | < 2e-16 |
|  | Family*Environment | 11 | 1286 | 117 | 6.567 | 8.27E-11 |
|  | Genotype*Environment | 1015 | 19244 | 19 | 1.065 | 0.131 |
|  | Residuals | 1650 | 30146 | 18 |  |  |
| NT | Environment | 1 | 934.5 | 934.5 | 889.235 | < 2e-16 |
|  | Family | 11 | 122.3 | 11.1 | 10.581 | < 2e-16 |
|  | Genotype | 1147 | 1344.9 | 1.2 | 1.116 | 0.0215 |
|  | Family*Environment | 11 | 101 | 9.2 | 8.734 | 3.22E-15 |
|  | Genotype*Environment | 1015 | 1316.9 | 1.3 | 1.235 | 8.22E-05 |
|  | Residuals | 1650 | 1733.9 | 1 |  |  |
| SGY | Environment | 2 | 618871 | 309435 | 892.123 | < 2e-16 |
|  | Family | 11 | 83674 | 7607 | 21.931 | < 2e-16 |
|  | Genotype | 1159 | 563910 | 487 | 1.403 | 1.73E-12 |
|  | Family*Environment | 19 | 53801 | 2832 | 8.164 | < 2e-16 |
|  | Genotype*Environment | 2040 | 992319 | 486 | 1.402 | < 2e-16 |
|  | Residuals | 2690 | 933034 | 347 |  |  |
| PGY | Environment | 2 | 10322143 | 5161072 | 133.9528 | < 2e-16 |
|  | Family | 11 | 9599721 | 872702 | 22.6505 | < 2e-16 |
|  | Genotype | 1159 | 62861091 | 54237 | 1.4077 | 1.00e-16 |
|  | Family*Environment | 19 | 5359413 | 282074 | 7.3211 | < 2e-16 |
|  | Genotype*Environment | 2040 | 102681425 | 50334 | 1.3064 | 4.87e-11 |
|  | Residuals | 2694 | 103797235 |  |  |  |
| TSW | Environment | 2 | 106473 | 53237 | 3634.438 | < 2e-16 |
|  | Family | 11 | 9204 | 837 | 57.12 | < 2e-16 |
|  | Genotype | 1159 | 31528 | 27 | 1.857 | < 2e-16 |
|  | Family*Environment | 19 | 3330 | 175 | 11.965 | < 2e-16 |
|  | Genotype*Environment | 2040 | 45577 | 22 | 1.525 | < 2e-16 |
|  | Residuals | 2690 | 39403 | 15 |  |  |

**Table S2**. Sorghum BCNAM trait means within each environment.

| Trait§ | Unit | Kobo | Meiso | Sheraro |
| --- | --- | --- | --- | --- |
| HE | cm | 1.29 | 2.91/6.15‡ |  |
| LS | Score (1-5) | 2.49 | 2.81 |  |
| NL | count | 8.76 | 9.95 | 13.60 |
| NT | count | 0.67 | 0.21 |  |
| SGY | gram | 62.83 | 54.64 | 38.20 |
| CPP | count | 8.45 | 9.08 | 15.67 |
| PGY | gram | 340.39 | 522.98 | 601.99 |
| TSW | gram | 29.48 | 20.80 | 19.67 |
| DF | days | 78.15 | 85.05 | 64.99 |
| DM | days | 127.04 | 122.66 | 90.22 |
| GFP | days | 48.89 | 37.61 | 25.22 |

§HE: head exsertion, LS: leaf senescence, NL: number of leaves, NT: number of tillers, SGY: single plant-based grain yield, CPP: count of plants per plot, PGY: plot-based grain yield, TSW: 1000-seed weight, DF: days to 50% flowering, DM: days to maturity, GFP: grain filling period, which was calculated by subtracting DM by DF.

‡Since HE at Meiso exhibited severe left-skewed distribution, Box-Cox transformation was conducted to transform original data to approximately normal distribution. 2.91 is the trait mean value obtained using Box-Cox transformed data, 6.15 is the trait mean value obtained using original data.
