## Supplementary figures and images for "Natural variation further increases resilience of sorghum bred for chronically drought-prone environments"

### Figure S1

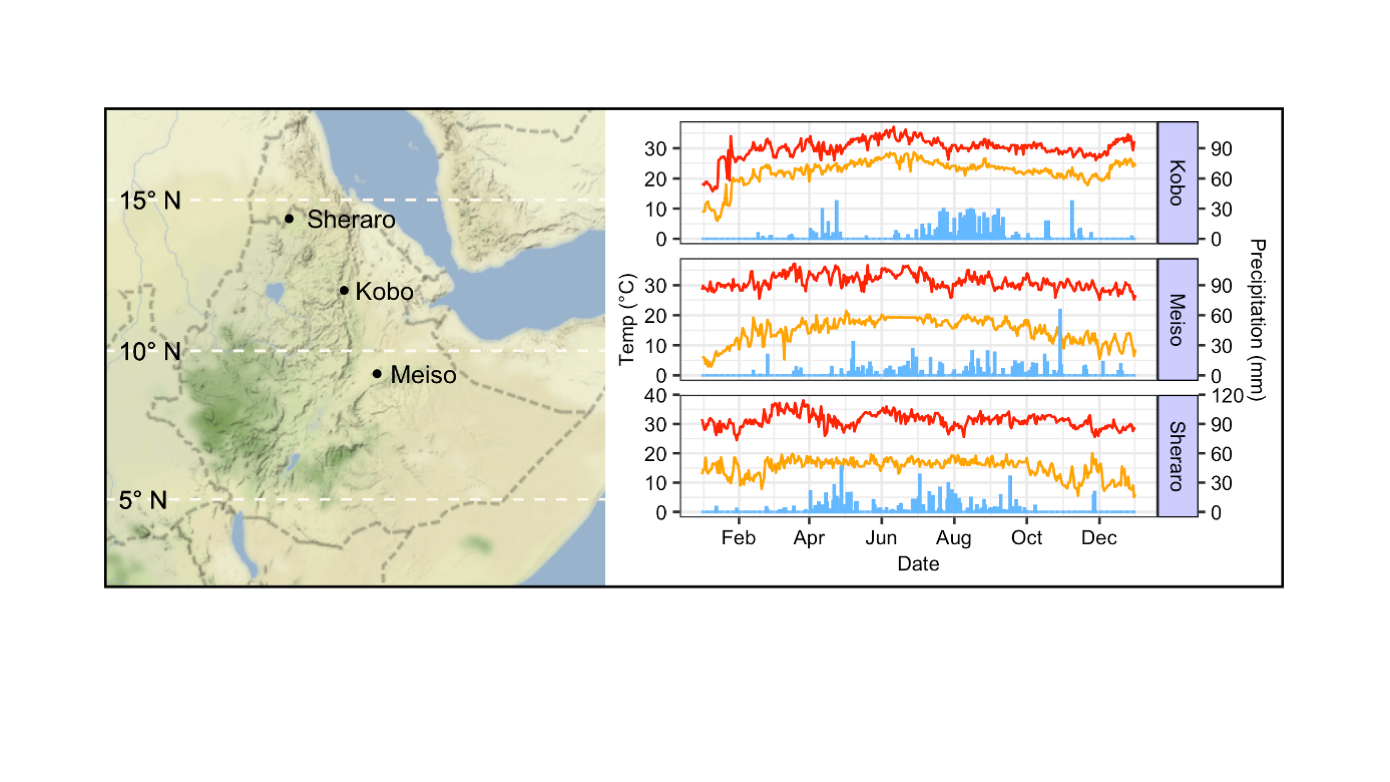

### Figure S2

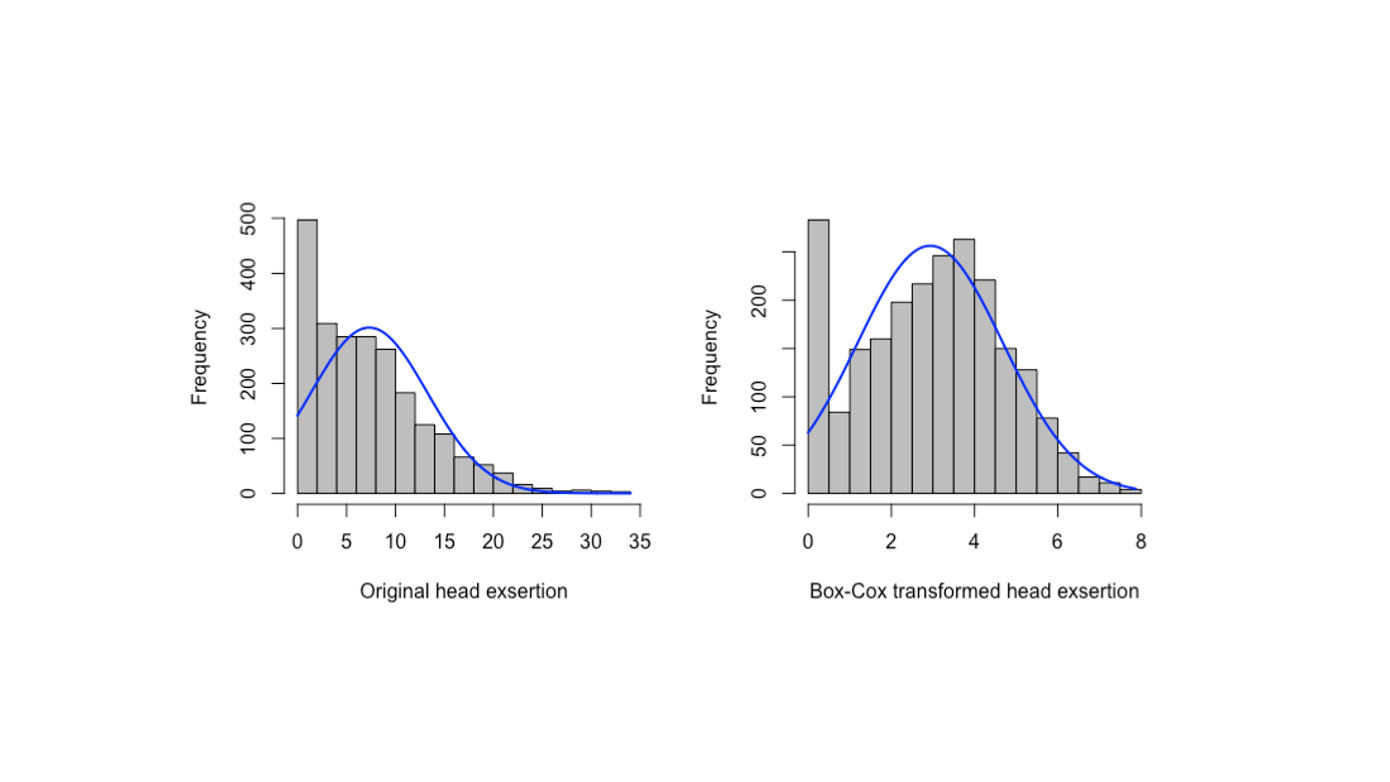

### Figure S3

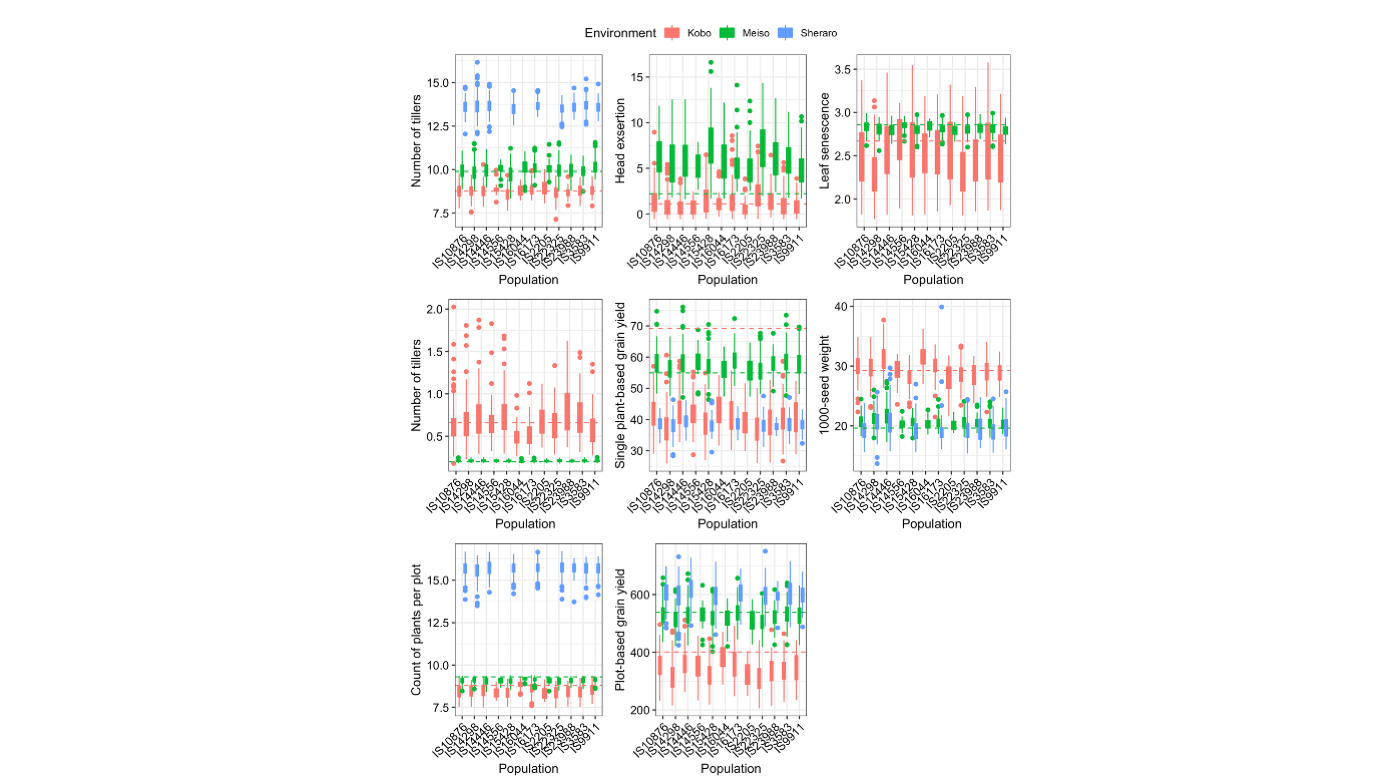

### Figure S4

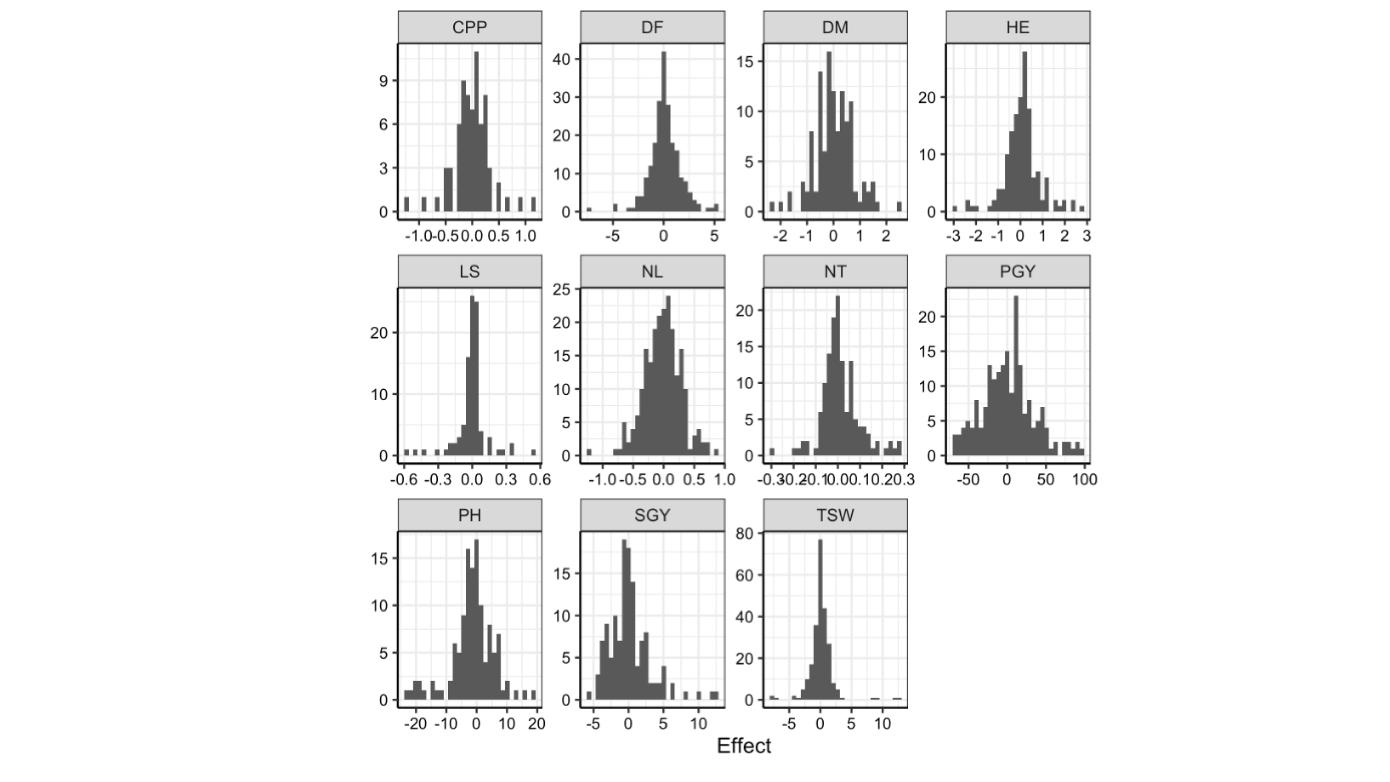
